# In-season internal and external training load quantification of an elite European soccer team

**DOI:** 10.1101/489187

**Authors:** Rafael Oliveira, João P. Brito, Alexandre Martins, Bruno Mendes, Francisco Calvete, Sandro Carriço, Daniel A. Marinho, Ricardo Ferraz, Mário C. Marques

**Affiliations:** Sports Science School of Rio Maior – Polytechnic Institute of Santarém, Portugal; Benfica Lab of Human Performance, Seixal, Portugal; Research Centre in Sport Sciences, Health Sciences and Human Development, Portugal; Research Centre on Quality of Life, Portugal; University of Beira Interior, Department of Sports Sciences, Covilhã, Portugal; Castelo Branco Football Association

**Author notes:** Corresponding Author: (RO).

**Keywords:** soccer training, internal load, external load, training load, periodization

## Abstract

Elite soccer teams that participate in European competitions often have a difficult schedule, involving weeks in which they play up to three matches, which leads to acute and transient subjective, biochemical, metabolic and physical disturbances in players over the subsequent hours and days. Inadequate time recovery between matches can expose players to the risk of training and competing whilst not fully recovered. Controlling the level of effort and fatigue of players to reach higher performances during the matches is therefore critical. Therefore, the aim of the current study was to provide the first report of seasonal internal and external training load (TL) that included Hooper Index (HI) scores in elite soccer players during an in-season period. Sixteen elite soccer players were sampled, using global position system, session rating of perceived exertion (s-RPE) and HI scores during the daily training sessions throughout the 2015-2016 in-season period. Data were analysed across ten mesocycles (M: 1 to 10) and collected according to the number of days prior to a match. Total daily distance covered was higher at the start (M1 and M3) compared to the final mesocycle (M10) of the season. M1 (5589m) reached a greater distance than M5 (4473m) (ES = 9.33 [12.70, 5.95]) and M10 (4545m) (ES = 9.84 [13.39, 6.29]). M3 (5691m) reached a greater distance than M5 (ES = 9.07 [12.36, 5.78]), M7 (ES = 6.13 [8.48, 3.79]) and M10 (ES = 9.37 [12.76, 5.98]). High-speed running distance was greater in M1 (227m), than M5 (92m) (ES = 27.95 [37.68, 18.22]) and M10 (138m) (ES = 8.46 [11.55, 5.37]). Interestingly, the s-RPE response was higher in M1 (331au) in comparison to the last mesocycle (M10, 239au). HI showed minor variations across mesocycles and in days prior to the match. Every day prior to a match, all internal and external TL variables expressed significant lower values to other days prior to a match (p<0.01). In general, there were no differences between player positions.

**Conclusions:** Our results reveal that despite the existence of some significant differences between mesocycles, there were minor changes across the in-season period for the internal and external TL variables used. Furthermore, it was observed that MD-1 presented a reduction of external TL (regardless of mesocycle) while internal TL variables did not have the same record during in-season match-day-minus.

## Introduction

Elite soccer teams that participate in European competitions have weekly schedules featuring up to three-matches that can lead to increased levels of fatigue, and higher risk of illness and injury [1]. The knowledge of internal and external training load (TL) helps coaches to design an effective individual and group training periodization in elite team sports [2-7] Djaoui et al., 2017; Jaspers et al., 2016; Malone et al., 2015; 2017; Nédélec et al., 2012; Stevens et al., 2017). However, it is only recently that some studies have described the in-season training periodization practices of elite football teams in more detail, including a comparison of training days within weekly microcycles [4, 7-9]. As an example, Malone et al. [4] found that a lowering TL in the last training day immediately before any given match differed from the other training days on several internal and external TL load variables such as session rated perceived exertion (s-RPE), plus total distance and average speed, respectively. In addition, some studies have shown limited variation through the in-season and have suggested that training in elite soccer has a regular load pattern [4, 5, 10, 11].

Moreover, several authors [1, 10, 12, 13] have claimed that it is also very important to monitor elite athletes’ health to provide further information concerning the details of player fatigue, stress, muscle soreness, need for recovery and sleep perception. These variables are commonly associated with biochemical (physical and physiological) and biomechanical stress responses, recognized as internal TL [13, 14]. On this issue, a valid and simple way to control internal TL is the session rating of perceived exertion (s-RPE) which showed correlations to the heart frequency training zones [15]. Furthermore, another way to quantity the level of fatigue, stress and delayed onset muscle soreness (DOMS) and the quality of sleep is the Hooper Index [12].

However, the simultaneous use of s-RPE and Hoper Index (HI) is limited. In fact, very few authors have studied the relationship between the use of the HI and s-RPE [10, 16]. Here, Clemente et al. [10] found a correlation between s-RPE and HI levels, and negative correlations between s-RPE and DOMS (p= −0.156), s-RPE and sleep (p= −0.109), s-RPE and fatigue (p= −0.225), ITL and stress (p= −0.188) and ITL and HI (p= −0.238) in 2-game weeks. On the other hand, Haddad et al. [16] failed to observe any association between HI and RPE. Therefore, further research is needed to clary this issue, specifically to validate these results during in-season. Subsequently, it is also necessary to quantify the external TL that is associated with the total amount of workload performed during training sessions and/or matches [13-14]. According to Halson [17] and Casamichana et al. [18], one easy and practical way to control training response for each player (e.g. frequency, time, total distance and distances of different exercise training intensity) is time-motion analysis by using a global positioning system (GPS).

Nowadays, researchers study the data collected during short training microcycles of 1-2-3 weeks [9-10, 13, 19], in mesocycles consisting of 4-10 weeks [20-22] and during longer training periods of 3-4 months [18, 23] and 10-month periods [11]. However, most of these studies have provided limited information regarding the TL, using only the duration and RPE without the inclusion of other internal and external TL variables such as HI or data collected from GPS. In addition, few studies [4-5, 10] have attempted to quantify TL with respect to changes between mesocycles and microcycles (both overall and between player’s positions) across an in-season.

Finally, the literature is somewhat inconclusive about establishing differences in TL for player positions not only amongst training sessions but also during the in-season across a full competitive season regarding training sessions, but there is information related to match-play data that reveals some differences for player positions [4, 24]. Therefore, the purpose of this study was twofold: a) quantify external TL in an elite professional European soccer team that played UEFA competitions across ten months of the in-season 2015/16 and b) quantify the internal TL using s-RPE and HI. For this purpose, we divided the in-season into ten months, following Morgan et al. [11], and used the match day minus approach used by Malone et al. [4] for data analysis. Additionally, we also compared player positions for both situations. We hypothesized that training load is lower on training days closer to the next match and that the intensities and volume remain constant throughout the competitive period.

## Materials and methods

### Participants

Nineteen elite soccer players with a mean ± SD age, height and mass of 26.3 ± 4.3 years, 183.5 ± 6.6 cm and 78.5 ± 6.8 kg, respectively, participated in this study. The players belong to a team that participated in UEFA competitions. The field positions of the players in the study consisted of four central defenders (CD), four wide defenders (WD), four central midfielders (CM), four wide midfielders (WM) and three strikers (ST). Inclusion criteria were regular participation in most of the training sessions (80% of weekly training sessions); the completion of at least 60 minutes in one match in the first half of the season and one match in the second half of the season. All participants were familiarised with the training protocols prior to the investigation. The study was conducted according to the requirements of the Declaration of Helsinki and was approved by the institution’s research ethics committee.

### Design

TL data were collected over a 39-week period of competition during the 2015-2016 annual season. The team used for data collection competed in four official competitions across the season, including UEFA Champion league, the national league and two more national cups from their own country, which often meant that the team played one, two or three matches per week. For the purposes of the present study, all the sessions carried out as the main team sessions were considered. This refers to training sessions in which both the starting and non-starting players trained together. In addition, all data collected from matches for the period chosen were considered. Only data from training sessions and matches were considered. Data from rehabilitation or additional training sessions of recuperation were excluded. This study did not influence or alter the training sessions in any way. Training data collection for this study was carried out at the soccer club’s outdoor training pitches. Total minutes of training sessions included warm-up, main phase and slow down phase plus stretching. Compensation minutes of matches were included in the collected data, however this measure is not revealed because the administration of the soccer club does not want to provide any information that could identify the team in this study.

### Methodology

The in-season phase was divided into 10 mesocycles or 10 months, respectively, as used by Morgans et al. [11] and because the coaches and staff of the club work by months. Training data were also analysed in relation to the number of days away from the competitive match fixture (i.e., match day minus). In a week with only one match, the team typically trained five days a week (match day [MD] minus [-]; MD-5; MD-4; MD-3; MD-2; MD-1), plus one day after the match (MD+1). This approach was used by Malone et al. [4].

#### External training load – training data

A portable global positioning system (GPS) units (Viper pod 2, STATSports, Belfast, UK) was used to monitor the physical activity of each player (external TL). This device provides position velocity and distance data at 10 Hz frequency. Each player wore the device inside a custom-made vest supplied by the manufacturer across the upper back between the left and right scapula. This position allows the GPS antenna to be exposed for a clear satellite reception. All players wore the same GPS device for each training session in order to avoid inter unit error [25]. Previously, some studies have been able to provide valid and reliable estimates of instantaneous and constant velocity movements during linear, multidirectional and soccer-specific activities by using this system [26, 27].

Following recommendations by Maddison & Ni Mhurchu [28], all devices were activated 30 minutes before data collection to allow the acquisition of satellite signals and synchronise the GPS clock with the satellite’s atomic clock. GPS data were then downloaded using the respective software package (Viper PSA software, STATSports, Belfast, UK) and were clipped to involve the main team session (i.e. the beginning of the warm up to the end of the last organised drill).

The metrics selected for the study were total duration of training session, total distance and high speed distance (HSD, above 19Km/h).

#### Internal training load – training data

Approximately 30 min before each training session, each player was asked to rate the perception of the quantity of fatigue, stress and DOMS and quality of sleep of the night that preceded the evaluation. The Hooper index scale of 1–7 was used, in which 1 is very, very low and 7 is very, very high (for stress, fatigue and DOMS levels) and 1 is very, very bad and 7 is very, very good (for sleep quality). The Hooper Index is the summation of the four subjective ratings [12].

Thirty minutes following the end of each training session, players were asked to provide an RPE rating, 0-10 scale [29]. Players were prompted for their RPE individually using a custom-designed application on a portable computer tablet. The player selected their RPE rating by touching the respective score on the tablet, which was then automatically saved under the player’s profile. This method helped minimise factors that may influence a player’s RPE rating, such as peer pressure and replicating other player’s ratings [30]. Each individual RPE value was multiplied by the session duration to generate a session-RPE (s-RPE) value [21, 31, 32].

### Statistical Analysis

Data were analysed using SPSS version 22.0 (SPSS Inc., Chicago, IL) for Windows statistical software package. Initially, descriptive statistics were used to describe and characterize the sample. Shapiro-Wilk and the Levene tests were used to assumption normality and homoscedasticity, respectively. ANOVA was used with repeated measures with Bonferroni post hoc, once variables obtained normal distribution (Shapiro-Wilk>0.05), to compare 10 mesocycles and to compare days away from the competitive match fixture. Results were significant in the interaction (p≤0.05). The effect-size (ES) statistic was calculated to determine the magnitude of effects by standardizing the coefficients according to the appropriate between-subjects standard deviation and was assessed using the following criteria: <0.2 = trivial, 0.2 to 0.6 = small effect, 0.6 to 1.2 = moderate effect, 1.2 to 2.0 = large effect and >2.0 = very large [33]. The associations between s-RPE and HI scores were tested with Spearman correlation. Data are represented as mean ± SD.

## Results

### In-Season Mesocycle Analysis (table 1)

For the duration of training sessions, M1 had more minutes than other mesocycles, especially M4 (ES = 6.77 [9.11, 4.44]), M5 (ES =9.64 [13.12, 6.16]), M6 (ES = 6.64 [9.14, 4.14]) which decreased, then increased some minutes to M7 and then decreased to M8 (ES = 6.17 [8.52, 3.82]), to M9 (ES = 5.83 [8.09, 3.59]) and to M10 (ES = 6.89 [9.47, 4.31]). M5 was the lowest, especially compared to M7 (ES = 5.72 [3.51, 7.93]) M8 (ES = 5.74 [3.53, 7.96]) and M10 (ES = 5.03 [3.03, 7.02]). There were no differences between player positions during the season (fig 1). For external load, total distance tended to decrease during the season. M1 and M3 saw a greater distance reached. M1 reached a greater distance than M5 (ES = 9.33 [12.70, 5.95]) and M10 (ES = 9.84 [13.39, 6.29]). M3 reached a greater distance than M5 (ES = 9.07 [12.36, 5.78]), M7 (ES = 6.13 [8.48, 3.79]) and M10 (ES = 9.37 [12.76, 5.98]). There were significant differences between player positions only in M1 for WD vs WM (ES = 4.87 [6.82, 2.92]), CM vs WM (ES = 5.07 [7.09, 3.06]) (fig 1); Average speed had few variations during the season. M3 reached the highest average speed while M10 reached the lowest (ES = 7.15 [9.81, 4.49]); high-speed running distance was higher in M1, especially compared to M5 (ES = 27.95 [37.68, 18.22]), which was the mesocycle with the lowest high-speed running, compared to M6 (ES = 5.89 [8.15, 3.63]), M7 (ES = 12.65 [17.15, 8.16]), M8 (ES = 6.31 [8.71, 3.92]), M9 (ES = 7.27 [9.97, 4.57]) and M10 (ES = 8.46 [11.55, 5.37]). There were significant differences between player positions only in M1 for CD vs WD (ES = 5.01 [3.02, 7.00]). For internal load, s-RPE was higher in M1, especially compared to M5 (ES = 5.17 [7.21, 3.13]) and M8 (ES = 3.87 [5.53, 2.21]), with a tendency to decrease until the end of the season to M10 (ES = 3.81 [5.46, 2.17]). There were no differences between player positions during the season (fig 1). HI had fewer variations during the season, reaching the highest value in M5 and the lowest value in M10 (ES = 3.47 [5.03, 1.92]). There were no significant differences between player positions.

**Fig 1.**
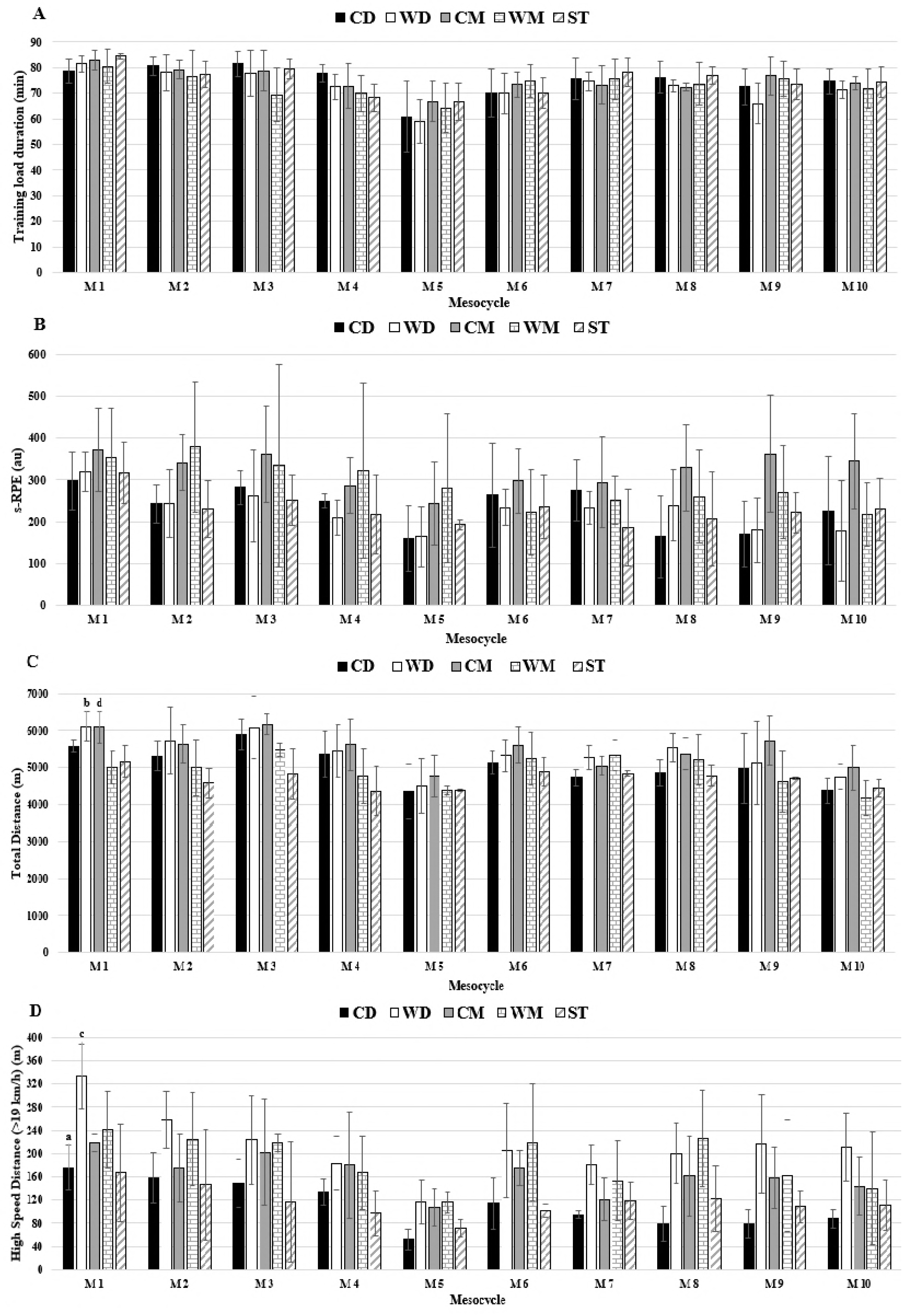
TL data for duration, s-RPE, total distance and HSD in respect to mesocycles between positions. Abbreviations: (A) duration; (B) s-RPE; (C) total distance; (D) HSD; (CD), central defenders; (WD), wide defenders; (CM), central midfielders; (WM), wide midfielders; (ST), strikers. a denotes significant difference in CD versus WD, (b) denotes significant difference in WD versus WM, (c) denotes significant difference in WD versus ST, (d) denotes significant difference CM versus WM, all P < 0.05.

There were associations between HI scores and s-RPE, HI scores and external TL variables, and S-RPE and external TL variables, but few correlations were found: stress and total distance in M2 (-6.34, p<0.01); fatigue and s-RPE in M9 (0.589, p<0.05); DOMS and s-RPE in M9 (0.487, p<0.05); fatigue and s-RPE in M11 (0.469, p<0.05); and HI total score and total distance in M11 (0.489, p<0.05).

**Table 1.** Training Load Data during the ten mesocycles for squad average, Mean ± SD

| Mesocycle<br>(number of<br>matches) | Duration (min) | Total Distance (m) | Average speed<br>(m/min) | HSD (m) | s-RPE (au) | HI (au) |
| --- | --- | --- | --- | --- | --- | --- |
| M1 (4) | 81.6±1.1<sup>c, d, e, g, h, i</sup> | 5589.1±100.1<sup>d, i</sup> | 68.6±1.1 | 227.0±13.7<sup>d, e, f, g, h, i</sup> | 331.9±21.6<sup>d, g, i</sup> | 11.9±0.8 |
| M2 (5) | 78.4±1.6<sup>d, i</sup> | 5248.2±156.2<sup>b, i</sup> | 66.8±0.9<sup>b</sup> | 192.3±17.0<sup>d, g</sup> | 287.3±22.6<sup>d</sup> | 12.1±0.8 |
| M3 (4) | 77.4±1.9<sup>d</sup> | 5691.4±132.1<sup>d, f, i</sup> | 74.0±1.7<sup>i</sup> | 181.9±18.9<sup>d</sup> | 298.4±33.2 | 11.7±0.7 |
| M4 (5) | 72.3±1.6 | 5111.4±173.9 | 70.7±2.2 | 152.2±15.4<sup>d</sup> | 256.9±26.6 | 12.6±0.7 |
| M5 (6) | 63.6±2.4<sup>f, g, i</sup> | 4473.5±136.4<sup>e, f</sup> | 71.0±2.1<sup>i</sup> | 92.3±6.6<sup>e, f, g</sup> | 208.6±25.9 | 13.0±0.7<sup>i</sup> |
| M6 (8) | 71.7±1.8 | 5231.8±123.0<sup>i</sup> | 73.2±1.7<sup>i</sup> | 162.9±15.3 | 250.5±22.1 | 11.4±0.9 |
| M7 (5) | 75.5±1.7 | 5041.9±70.5<sup>i</sup> | 67.2±1.9 | 133.6±10.3 | 247.8±20.4 | 11.6±1.1 |
| M8 (4) | 74.5±1.2 | 5149.5±112.5<sup>i</sup> | 69.3±1.3<sup>i</sup> | 157.8±15.4 | 239.8±25.8 | 10.6±0.8 |
| M9 (7) | 72.9±1.8 | 5026.7±204.1 | 69.0±2.1<sup>i</sup> | 144.8±15.9 | 240.8±25.5 | 10.8±0.8 |
| M10 (4) | 73.3±1.3 | 4545.4±111.7 | 62.2±1.6 | 138.5±14.7 | 239.3±26.7 | 10.2±0.9 |

M= mesocycle (1, 2, 3, etc.); min= minutes; m=meters; HSD = high-speed running distance; s-RPE= session rating of perceived effort; HI = Hooper index; au=arbitrary units. a denotes difference from M2, b denotes difference from M3, c denotes difference from M4, d denotes difference from M5, e denotes difference from M6, f denotes difference from M7, g denotes difference from M8, h denotes difference from M9, i denotes difference from M10, all P < 0.05

### In-Season Match-Day-Minus Training Comparison (table 2)

For duration of training sessions, MD-5 was higher than MD-4 (ES = 4.44 [6.27, 2.62]), MD-3 (ES = 5.69 [7.90, 3.49]), MD-2 (ES = 6.49 [8.94, 4.03]) and MD+1 (ES = 42.61 [57.4, 27.81]), with the exception of MD-1 (ES = -6.34 [-3.94, -8.75]). MD-4 (ES = -4.44 [-6.27, -2.62]), (ES = 42.61 [57.4, 27.81]), MD-3 (ES = -13.14 [-8.48, -17.79]), (ES = 37.33 [50.31, 24.36]) and MD-2 (ES = -18.24 [-11.85, -24.64]) (ES = 43.92 [59.17, 28.67]) were higher than MD-1 and MD+1, respectively. MD-1 was the highest and MD+1 was the lowest (ES = 61.40 [82.70, 40.10]). No differences were found between players positions (fig 1). For external load, total distance was higher in MD-5 than MD-4 (10.73 [14.58, 6.86]), MD-3 (8.88 [12.11, 5.65]), MD-2 (16.06 [21.71, 10.41]), MD-1 (30.47 [41.07, 19.87]) and MD+1 (16.23 [21.93, 10.52]). MD-4 was higher than MD-2 (6.31[8.70, 3.91]), MD-1 (28.24 [38.07, 18.40]) and MD+1 (9.09 [12.39, 5.79]). MD-3 was also higher than MD-2 (9.30 [12.67, 5.93]), MD-1 (17.51 [23.66, 11.37]) and MD+1 (10.80 [14.67, 6.93]). MD-2 was higher than MD-1 (32.04 [43.18, 20.89]) and MD+1 (6.03 [8.34, 3.73]), and MD+1 was higher than MD-1 (7.42 [4.67, 10.17]). There were significant differences in MD-2 between WD vs ST (5.13 [9.19, 1.07]) and CM vs ST (5.01 [9.01, 1.02]). Average speed was higher in MD-5 than MD-4 (6.01 [8.29, 4.71]), MD-3 (3.81 [5.45, 2.16]), MD-2 (9.20 [12.54, 5.87]), MD-1 (24.36 [32.86, 15.86]) and MD+1 (-12.69 [-8.19, -17.20]). MD-4 was higher than MD-2 (6.37 [8.79, -3.96]), MD-1 (41.11 [55.39, 26.83]) and MD+1(-14.05 [-9.09, -19.02]). MD-3 was also higher than MD-2 (10.56 [14.35, 6.77]), MD-1 (46.36 [58.42, 28.31]) and MD+1(-13.63 [-8.80, -18.45]). MD-2 was higher than MD-1 (45.96 [61.92, 30.01]) and MD+1 (-14.63 [-9.47, -19.80]), and MD+1 was higher than MD-1 (17.44 [11.32, 25.56]). No differences were found between player positions (fig 2); high-speed running distance was higher in MD-5 than MD-2 (4.22 [5.98, 2.46]), MD-1 (10.75 [14.61, 6.90]) and MD+1 (7.05 [9.67, 4.42]). MD-4 was higher than MD-2 (2.33 [4.01, 1.06]), MD-1 (14.49 [19.60, 9.37]), MD+1 (7.71 [10.55, -.86]). MD-3 was also higher than MD-2 (2.35 [3.62, 1.08]), MD-1 (14.04 [19.00, 9.08]) and MD+1 (6.41 [8.85, 3.99]). MD-2 was higher than MD-1 (13.37 [18.11, 8.64]), MD+1 (4.89 [6.85, 2.94]) and MD+1 was higher than MD-1 (3.44 [1.89, 4.98]). In MD-3 there were significant differences between player positions (fig 2) for CB vs WD (4.94 [1.01, 8.89]). In MD-2 there were significant differences between player positions CD vs WD (7.81 [2.05, 13.57]), CD vs WM (5.74 [1.31, 10.17]) and WD vs ST (6.02 [10.62, 1.41]). In MD-1 there were significant differences between player positions CD vs WD (4.93 [0.99, 8.86]) and WD vs ST (5.03 [1.03, 9.04]). For internal load, s-RPE was higher in MD-3 than MD-2 (2.81 [4.19, 1.42]), MD-1 (6.20 [8.56, 3.84]) and MD+1 (17.08 [23.08, 11.08]). MD-5 was higher than MD-1 (5.42 [7.54, 3.30]) and MD+1 (15.47 [20.92, 10.02]). MD-4 was higher than MD-2 (2.45 [3.75, 1.15]), MD-1 (5.74 [7.95, 3.52]) and MD+1 (9.77 [13.30, 6.25]). No differences were found between player position (fig 1). HI had few variations during the MD minus with the exception of MD+1, which were higher than MD-5 (7.43 [10.18, 4.67]), MD-4 (6.60 [9.08, 4.11]), MD-3 (6.60 [9.08, 4.11]), MD-2 (6.29 [8.68, 3.90]) and MD-1 (6.90 [9.49, 4.32]). No differences were found between player positions (fig 2).

**Table 2.**
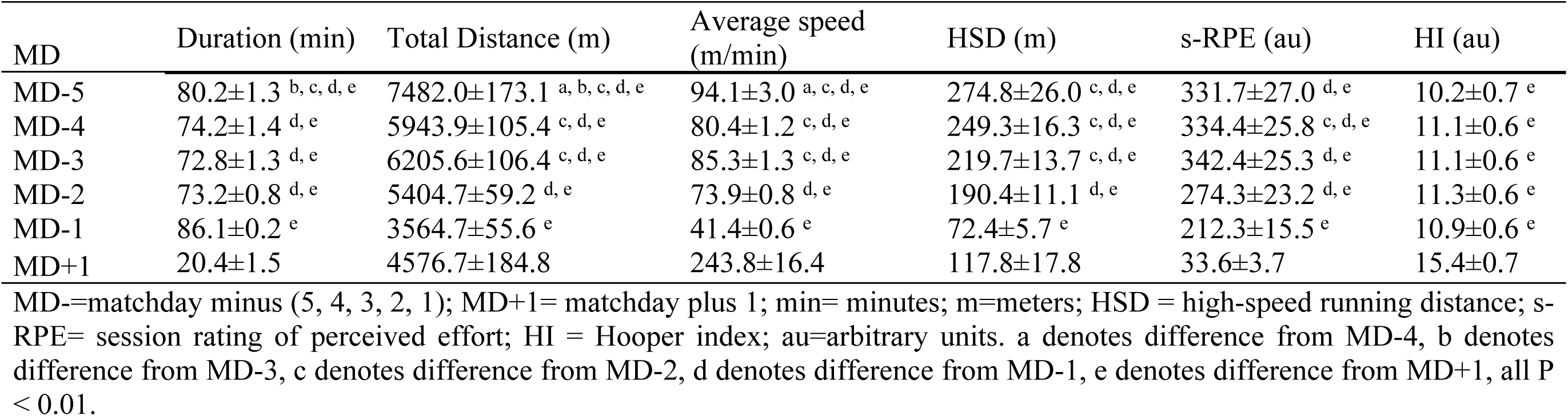
Training Load Data during the MD minus for squad average, Mean ± SD

**Fig 2.**
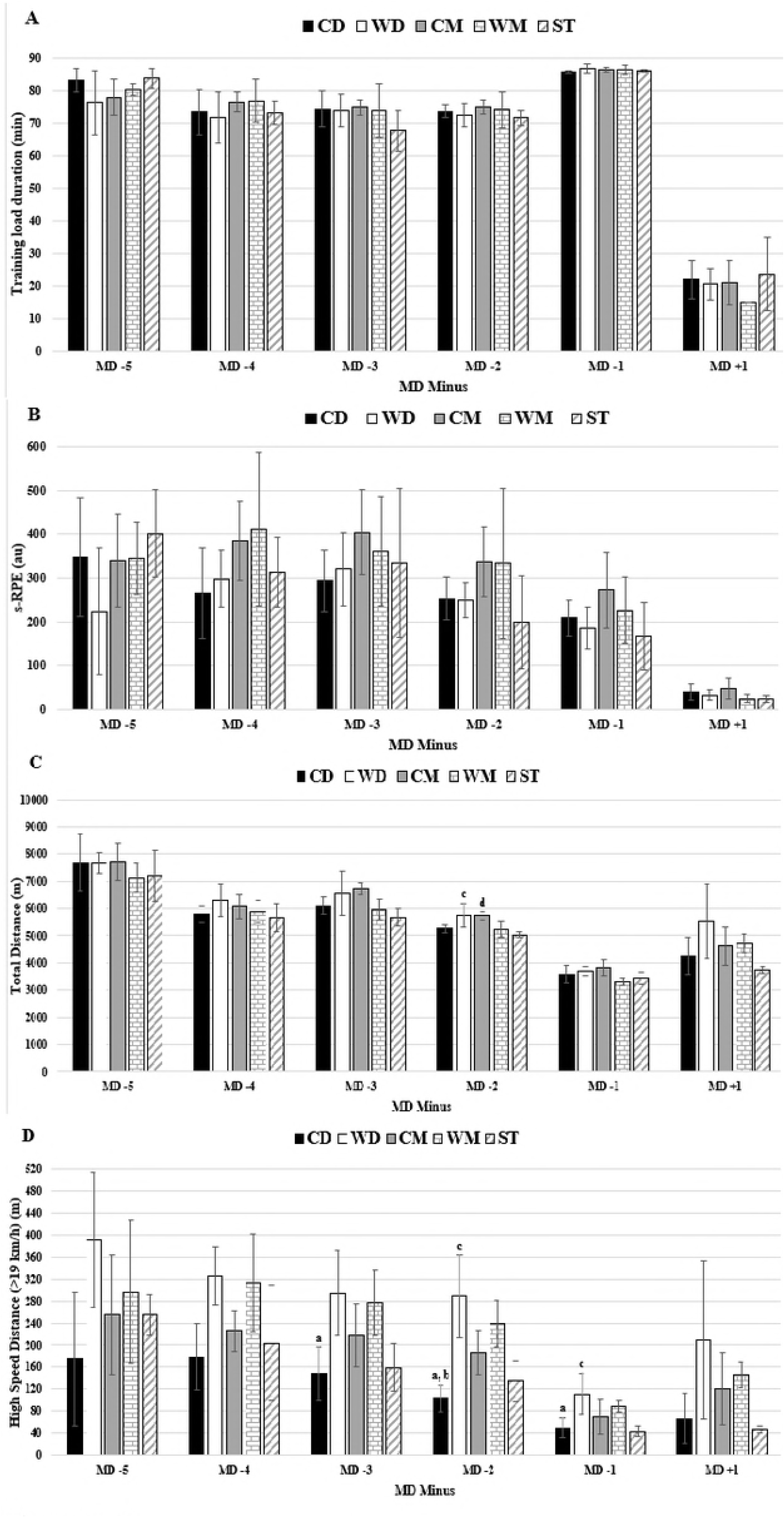
TL data for duration, s-RPE, total distance and HSD in respect to days before a competitive match between positions. Abbreviations: A) duration; (B) s-RPE; (C) total distance; (D) HSD; (CD), central defenders; (WD), wide defenders; (CM), central midfielders; (WM), wide midfielders; (ST), strikers. (a) denotes significant difference in CD versus WD, (b) denotes.

## Discussion

The purpose of the present study was to quantify the internal and external TL carried out by an elite soccer team during the in-season (10 mesocycles).

### In-season mesocycle analysis

For external TL variables, it was observed that the players covered a greater total distance at the start (M1 and M3) compared to the final mesocycle (M10) of the in-season, with an estimated difference of 1044m and 1146m, respectively. The higher distances covered at the beginning of the in-season may be due to the coaches still having some emphasis on physical conditioning immediately after the pre-season. In addition, the lower values in distance covered for M10 could be associated with the in-season ending and consequently a reduction in external TL.

According to Impellizzeri et al. [21] and Alexiou & Coutts [23], the competitive matches represent the greatest TL that soccer players typically experience. In addition, Malone et al. [4] and Los Arcos et al. [34] reported that total distance values were significantly higher at the start of the annual in-season compared to the final stage 1304 (434 – 2174) m, ES = 0.84 (0.28 – 1.39) and (ES = from – 0.56 to -1.20), respectively. These previous data corroborate our results because it was possible to observe higher values in M1 compared to M10, although M5 had the lowest values for total distance (table 1).

The present data suggest that in-season variability in TL is very limited and only minor decrements in TL across the in-season might occur. Apparently, this TL maintenance during the in-season could be associated with the importance of the recovery activities after the matches and the decisions made to reduce TL until the next match [35]. Furthermore, elite European soccer teams training programmes remain constant during all mesocycles of the in-season and corroborate the suggestion made by Malone et al. [4] because there is a need to win matches that does not allow the reaching of a specific peak for strength and conditioning.

The average total distance covered was 5111m (4473-5691m) which was similar to the 5181m value reported by Malone et al. [4] and slightly higher than those reported by Gaudino et al. [20] (3618-4133m). However, both the distances covered in the present study and in Gaudino et al. [20] study fell short in comparison to those reported by Owen et al. [19] (6871m) because their study only included data from training sessions. This means that the study conducted by Owen et al. [19] reported higher distances covered even with lower training sessions. In terms of high-speed distance, the values (average 118m) fall within the range of that of Gaudino et al. [20] (88–137m) across different positions.

The results indicate that TL variables demonstrated limited relevant variation between player positions (see fig 1 and 2). Competitive matches have been quantified as the most demanding session (i.e. greatest TL) of the week [7, 24, 34, 36]. For instance, Di Salvo et al. [37] reported that CM generally cover more distances compared to other positions during competitive matches. This result corroborates the current results because CM (5502m) covered more total distance than CD (5052m), WD (5388m), WM (4918m) or ST (4694m), but without statistical significance. In addition, when we compared the distance covered in high-speed running zones (zones 4+5) during in-season mesocycle analysis to positions played, a significant difference was found between positions only for M1 when comparing CD vs WD and WD vs WM. There was no other difference between player positions in all mesocycles (fig 1). These results suggest that the WD (212.7m) and WM (186,8m) positions resulted in higher effort (>19km/h) during training when compared to all other positions (CD=112.2, CM=164.1, ST=116.1m). Further, every position saw similar efforts at low speed distance (CD=4563.7; WD=4724.5, CM=4767.8, WM=4340.4, ST=4233.3m) which is in opposition to other studies [24, 37, 38].

Regarding internal TL, the s-RPE response was higher in M1 (331au) in comparison to the last mesocycle (M10, 239au) which is in line with data from external TL total distance and HSD variables. However, it was found that in the middle of the season (M5) there was a lower response (208au) for this parameter. This finding could be associated with some interruption for TL carried out during training sessions due to the Christmas period and with an increase in the number of matches played in M5 (6 matches). In general, there were no differences between player positions (see fig 1). Therefore, it appears that there is no marked variation in internal TL across 10 mesocycles during the in-season. Some studies [4, 5, 10, 11] have also reported the limited relevant variation in TL across the in-season. This seems to suggest that professional soccer daily training practices follow a regular load pattern because they are linked to higher congestive periods of matches. Furthermore, the importance of the recovery activities following matches and the decisions made to reduce TL between matches to prevent fatigue during this period can also play an important role in this constant TL [35].

Moreover, the data provides relevant information to quantify internal TL, measured by s-RPE during microcycles and mesocycles. This may provide relevant information to establish guidelines for soccer training periodization. The average of s-RPE during microcycles TL was 254.8au (range 33-342au). These values are lower than those reported by Scott et al. [22] (297au: range 38-936au), but similar to Jeong et al. [39] study: 174-365au. for elite soccer players. The difference between our study and the experiment conducted by Jeong et al. [39] could be attributed to the fact that they used a sample of Korean professional players rather than top elite European soccer athletes that competed in European competitions. The s-RPE values were also lower than the 462au of semi-professional soccer players reported by Casamichana & Castellano [18]. Another explanation for the lower values could be related to the amount of matches during each week and amongst mesocycles. It should be reemphasised that we studied a top-class elite professional European soccer team. The range of s-RPE for mesocycles of the in-season was 208-331au. Overall it would appear that in comparison to top elite soccer players, the internal TL employed by our study falls within the boundaries of what has been previously observed [18, 22, 39].

Haddad et al. [16] suggested that s-RPE is not sensitive to the subjective perception of fatigue, DOMS or stress levels [16]. In contrast, however, Clemente et al. [10] stated that s-RPE could be a reliable tool to quantify the internal TL and therefore could be a good indicator for coaches and for practical applications in team sports training. Data presented in the current experiment seems to corroborate this statement, indicating that s-RPE can be an effective tool to measure training intensity in elite European soccer teams. On this subject, some studies have stated that RPE may be a physiological and volatile construct that could be different according to the cognitive focus of the player [40-42]. Nevertheless, Renfree et al. [43] reported that RPE can be dissociated from the physiological process through a variety of psychological mechanisms. Therefore, RPE could be an oversimplification of the psychophysiological perceived exertion and a non-conclusive measure for capturing a wide range of sensations experience [40, 41, 43]. Another major point is that RPE was collected 30 min after the end of each training session and it would be pertinent to check if there is some variation during the training session, as contended by Ferraz et al. [41]. These arguments may justify the fact that there were no differences in s-RPE between training days as well as the absence of a relationship with the external TL results.

Comparing player positions, there were no differences for HI scores; this was not supported by Clemente et al. [10] although their study was based on data from one vs two-matches week (p< 0.05). To the best of our knowledge, this is the first study to analyse HI scores during an entire in-season. Clemente et al. [10] showed that central defenders (12.46 ± 2.54) and wide midfielder (12.42 ± 3.44) had higher values of HI scores than strikers (12.18 ± 4.84) and wide defenders (12.16 ± 3.04). Centre midfielders had the lowest HI scores (10.34 ± 3.87). Despite these, the authors found several significant differences between positions but, in general, these values were small. A possible explanation for these non-consensual results could be associated with the differences in soccer TL.

In soccer training, due to the extensive use of small-sided matches and the different physical (e.g. running) requirements associated with each position [37, 44, 45], training demands can be markedly different between individuals [13, 46, 47]. This hypothetical difference in TL could be amplified considering that only 11 players can start each official match, and therefore a considerable number of players per team are not exposed to the TL of the match.

As suggested by Clemente et al. [10] study, we also correlated HI scores with s-RPE and external TL variables, and some correlations could be observed: stress and total distance in M2 (-6.34, p<0.01); fatigue and s-RPE in M9 (0.589, p<0.05); DOMS and s-RPE in M9 (0.487, p<0.05); fatigue and s-RPE in M11 (0.469, p<0.05); and HI total score and total distance in M11 (0.489, p<0.05). These results are not in line with the literature, which suggests non-significant correlations (r=0.20) between s-RPE and perceived quality of sleep (from the Hooper questionnaire) [10, 48]. However, Thorpe et al. [49] reported associations between s-RPE and perceived fatigue, but not with perceived quality of sleep. It is important to note that this last study analysed data for short periods of training (microcycles). Therefore, since our study also comprised longer periods of training, we can assume that this could have influenced the current results.

### In-season match-day-minus training comparison

In the present study, we also investigated the TL pattern in respect to number of days prior to a match during the in-season phase.

For external TL, our data provided the following pattern by decreasing values from until MD-1: MD-5 > MD-4 < MD-3 > MD-2 > MD-1 for total distance and average speed, MD-5 > MD-4 > MD-3 > MD-2 > MD-1 for HSD (table 2). Our results are in line with elite English Premier League players for total distance and average speed, remaining similar across all days except for MD-1 in which the load was significantly reduced [4].

We also observed a noticeable consistent variation in external TL, total distance covered, in MD-1 when the load was significantly reduced in comparison with the rest of the training days. Our data corroborates with some studies [4, 8, 49].

Finally, MD+1 revealed significant result despite the limited training duration (~20 min). The average speed and HSD has higher values than all other match days minus. One argument that can justify these results could be the high-intensity applied by the coach (which was not controlled in this study). Another explanation is related to the context, competitive schedule and the objectives defined for TL management, once MD+1 had little duration (20min). Another possible justification could be associated with a training session of recuperation with lower load for starters and a “normal” training session for non-starters.

When we compared HSD (above 19Km/h) during in-season match-day-minus by positions, a significant difference was found between positions when comparing WD vs ST and CD vs WD, CD vs WM in MD-2 in MD-2. In addition, when we compared total distance covered, a significant difference could be observed between CD (149m) vs WD (295m) in MD-3, CD (103m) vs WD (289m) in MD-2 and CD (49m) vs WD (111m) in MD-1; CD (103m) vs WM (240m), WD (289m) vs ST (134m) in MD-2; and also WD (111m) vs ST (43m) in MD-1 (fig 2). These results are in line with other studies [24, 37-38] that reported that CM players have consistently been found to cover more distance in general while WM players cover more distances at high-intensity running speed.

Regarding match days, Reilly & Thomas [50] and Rienzi et al. [51] stated that higher distances are covered by midfield players (11.5km); however, Bangsbo [52] reported that elite defenders and strikers covered approximately the same distance (10-10.5km). This may be due to the nature and role of the position inside the team, as well as coaching strategy and/or game plan. During training sessions, the coach or the conditioning staff may find it advantageous to model training to elicit similar effort or experience the same training load regardless of position.

For internal TL, s-RPE data presented a non-perfect pattern by decreasing values from until MD-1: MD-5 < MD-4 < MD-3 > MD-2 > MD-1 for s-RPE (table 2), but none between player positions (fig 2). We also observed a noticeable consistent variation in s-RPE on MD-1 in elite soccer players, when the load was significantly reduced in comparison with the rest of the training days [4, 8, 49]. In addition, the data presented by s-RPE is associated with external TL variation.

Furthermore, HI scores revealed no variation in days prior to the match. These results are in line with those reported by Haddad et al. [16], where it was suggested that fatigue, stress, DOMS and sleep are not major contributors of perceived exertion during traditional soccer training without excessive TL. Our results also do not support Hooper and Mackinnon [12] study because self-reported ranking of well-being does not allow the provision of efficient mean of monitoring internal TL.

In opposition to the results presented for external in MD+1, internal TL, s-RPE has a lower value than all other match days (33.6 au) but HI has a higher value than all other match days (15au) (table 1). These results are associated with an accumulative high-intensity training session between MD-5 and MD-2

#### Practical Applications and Limitations

This study provides useful information relating to the TL employed by an elite European soccer team that played in a European Competition. It provides further evidence of the value of using the combination of different measures of TL to fully evaluate the patterns observed across the in-season. For coaches and practitioners, the study generates reference values for elite players which can be considered when planning training sessions. However, it is important to remember that the in-season match-day-minus training comparison was analysed by mean values and microcycles/weeks (7-day period) of the in-season have different patterns, as mentioned before. Another limitation is related to the numerous true data points missing across the 39-week data collection period due to several external factors beyond our control (e.g. technical issues with equipment, player injuries, and player transfers).

## Conclusions

In summary, we provide the first report across 10 mesocycles of an in-season that included HI scores and s-RPE to measure internal TL plus distances covered at different intensities measured by GPS, in elite soccer players that played European competitions. Our results reveal that although there are some significant differences between mesocycles, there was minor variation across the season for the internal and external TL variables used. In addition, it was observed that MD-1 presented a reduction of external TL during in-season match-day-minus training comparison (regardless of mesocycle) (i.e. reduction of total distance; five different training intensity zones) and internal TL (s-RPE). However, the internal TL variable, HI did not change, except for MD+1. This study also provided ranges of values for different external and internal variables that can be used for other elite teams.

## Acknowledgements

The authors would like to thank the team’s coaches and players for their cooperation during all data collection procedures.

This project was supported by the National Funds through FCT—Portuguese Foundation for Science and Technology (UID/DTP/04045/2013)—and the European Fund for Regional Development (FEDER) allocated by European Union through the COMPETE 2020 Programme (POCI-01-0145-FEDER-006969)—competitiveness and internationalization (POCI). The authors disclose funding received for this work from any of the following organizations: National Institutes of Health (NIH); Welcome Trust; Howard Hughes Medical Institute (HHMI); and other(s).

## Author Contributions

**Conceptualization:** RO, JB, RF, MCM.

**Data curation:** BM, FC, SC.

**Formal analysis:** RO, JB.

**Funding acquisition:** DM, RF, MCM.

**Investigation:** RO, JB.

**Methodology:** RO, JB, RF, MCM.

**Project administration:** RO, JB, BM, FC, SC.

**Resources:** RO, JB, AM, DM, RF, MCM.

**Software:** RO, JB.

**Supervision:** RO, JB, RF, MCM.

**Visualization:** RO, JB, RF, MCM.

**Writing – original draft:** RO, JB, AM, RF, MCM.

**Writing – review & editing:** RO, JB, RF, MCM.

